# The Impact of Body Mass Index on Physical Activity and Cardiac Workload

**DOI:** 10.1101/2021.09.26.461877

**Authors:** Myra Jane Bloom, Lakin Mckenzie Brown, Scarlet Rae Jost, Andrew Stuart Ian Donald Lang, Nancy Viola Mankin, Zachary William Mast, Ericka Rachel McMahan, Jonathan Abdou Merheb, Philip Paul Nelson, Joshua Chinweoke Nnaji, Enrique Francisco Valderrama

## Abstract

**Background:** Having an abnormal body mass index (BMI) adversely affects cardiac workload and level of physical activity.

**Objective:** To examine the relationship between cardiac workload, physical activity, Sex, and BMI.

**Methods:** The number of steps taken per day (Steps) and minutes per week spent in targeted heart rate zones were collected from primarily first and second year university students (n = 1,801; 62% female) over a standard, 15-week long semester. Other data collected included BMI, Sex, Age, and Class Standing. Sex differences in BMI, Steps, and training heart rate zone (heart rates above 50% of max) minutes (THR) were evaluated, correlations between the study parameters were analyzed, and one-way ANOVA was used to test between competing models. The values p < .05 were considered statistically significant.

**Results:** Statistically significant (p < .05) differences between males and females were found for Steps, THR, and BMI. Males were more physically active but spent 18% less time with heart rates above 50% of max. Students who had abnormal BMI values, both low and high, experienced greater cardiac workload (p < .05), even though they were found to be less physically active (p < .05).

**Conclusion:** Our study revealed that university students with abnormal BMI values experienced greater cardiac workload, even though they are less physically active. Thus, physical fitness and healthy lifestyle interventions should also include underweight students in addition to students who are overweight or obese.

## Introduction

Obesity in the United States has reached epidemic proportions with 42.4% of the adult population being obese^1^ and nearly 20% of children and adolescents being obese.^2^ For college students, recent studies have found declining levels of fitness combined with an increase in percent body fat for both males and females.^3-6^ Starting as freshmen, students gain weight at a significant level throughout their four years of college.^4,7,8^ A closer look at body mass index (BMI) and physical activity (PA) levels of college students and their relationship to cardiovascular health is needed to better understand how to best support students’ health and wellbeing in college.

Using BMI to measure body fat has both strengths and limitations. The strength is its simplicity of calculation from just weight and height^9^ which does not require specialized or expensive equipment that other more thorough laboratory methods use.^10^ Conversely, BMI fails to account for actual body composition, i.e. the difference between lean muscle and body fat.^11^ For example, for adults with the same BMI value, women typically have approximately 10% higher percent body fat than men.^12,13^

Some studies have also suggested that the cut-off points for different BMI categories should be altered for different populations.^14^ Several alternative measures of body fat include dual energy x-ray absorptiometry (DEXA), underwater weighing, waist circumference, waist-hip ratio, and analysis of skinfold thickness.^11^ However, these alternative measurements can be expensive, intrusive, or may need to be studied further. Therefore, BMI remains a commonly used body composition-based measure of health.

Research has shown that BMI is correlated with an increased risk of coronary heart disease.^15^ Additionally, higher resting heart rate (RHR) increases one’s risk for cardiovascular disease (CVD) and positively correlates with indicators of unhealthy lifestyles (e.g., higher blood pressure, abnormal BMI, high total cholesterol, and physical inactivity^16^). Underweight individuals with a BMI less than 18.5 kg**·**m^-2^ tend to be overlooked in relation to diagnosing risk factors for CVD. Most major studies have only looked at increased BMI (i.e., overweight and obese) as a risk factor for CVD, but have failed to understand risk factors associated with being underweight. For instance, Cooney^16^ found that a higher RHR, associated with higher BMI values, increased one’s risk for cardiovascular disease. However, Park et al.^17^ analyzed the results of BMI as a risk factor for CVD and found that underweight individuals had a 19.7% increased risk for CVD when compared with normal-weight individuals (n = 491,773). Comparatively, however, overweight (BMI between 25 kg**·**m^-2^ and 30 kg**·**m^-2^) individuals had a 50% greater risk and obese (BMI > 30 kg**·**m^-2^) individuals had a 96% greater risk for CVD. While being underweight has less of a risk for CVD than being overweight or obese, there is still an increased risk when compared to normal-weight individuals, especially for individuals under the age of 40. As such, when doing a comparative health study that includes BMI, the underweight category of BMI needs to be considered, especially for individuals under the age of 40.

In a meta-analysis of 40 articles investigating the relationship between BMI and physical activity, Jakicic et al.^18^ found a strong relationship between physical activity and prevention of weight gain in young adults. Young adults, who were found to get at least 150 minutes per week of moderate intensity exercise, showed less weight gain than those who get less than 150 minutes per week. For 20-29 year olds, Frank et al.^19^ found that, when compared with no activity, those who exercised 150 minutes per week (based on objective, activity data) had lower BMI and total cholesterol and those who self-reported exercising 150 minutes per week (subjective data) had higher HDL (“good” cholesterol) and lower diastolic blood pressure (n = 5,395, 47.9% female). As such, Frank et al.^19^ recommended that people aged 20-29 increase their PA levels. Similarly, individuals who are less physically active had higher maximum heart rates than those who were more active, indicative of increased cardiac workload.^20^ Cho, Lee, & Kim^21^ compared PA and sedentary behaviors with the BMI of Korean adolescent students. Using World Health Organization Asia-Pacific (AP) standards of obesity, it was found that significantly fewer students with a BMI of 18.5 kg**·**m^-2^ or lower (AP underweight) or 23.0 kg**·**m^-2^ and higher (AP overweight and obese) met the PA requirements than the students with a BMI between 18.5 kg**·**m^-2^ and 23.0 kg**·**m^-2^ (AP normal).

Davis et al.^22^ and Kelishadi et al.^23^ found that there was no significant sex difference in BMI for high school students, while Olfert^24^ found that college freshman males have slightly higher BMI values than college freshman females (23.4 kg**·**m^-2^ as compared to 23.8 kg**·**m^-2^, p = .0516). Those with metabolic disorders, often associated with abnormal RHR, showed elevated rates of BMI, cholesterol levels, glucose levels, and triglycerides.^25^ Furthermore, increased exercise has been shown to reduce CVD risk.^26^

Numerous studies have explored the validity of Fitbit in its measurement of HR and steps^27-29^ along with its effectiveness of measuring sleep patterns.^30^ Diaz et al.^31^ found a high correlation (0.77 - 0.85) for wrist devices when compared to manually counting steps. These studies have shown the reliability of data collected from Fitbit devices. Additionally, there is a lack of research that incorporates Fitbit-derived data among college students and its relationship to BMI, heart rate, and sex. From the current study, a better understanding of the relationship between BMI, physical activity, and cardiac workload will provide more effective strategies to increase the health of university students.

Quer et al.^30^ found a non-linear U-shaped relationship between RHR and BMI using data provided by Fitbit (n = 92,457 adults). As a person’s BMI was farther from the range that is considered normal (i.e., 18.5 kg**·**m^-2^ – 25 kg**·**m^-2^), their RHR would be higher. The current study will explore if this relationship exists amongst college students. In addition to HR measurements, PA will also be measured. Our study uses Fitbit-collected data to examine the relationships between BMI, Training Heart Rate zone (heart rates above 50% of max) minutes (THR), Sex, and PA of college students. The objective of the present study was to find significant correlations that would improve the understanding of BMI-related health and wellbeing factors for college students.

## Methods

### Participants

The data in this study were collected from 1,801 primarily first- and second-year students enrolled in various health and physical exercise (HPE) courses required as part of our institution’s whole person education curriculum. The sample was one of convenience as all students at our institution are enrolled in our HPE program and only a handful decided not to opt in to the study. All study participants gave consent to the University to collect daily step counts and HR data automatically from Fitbit devices. All data was then synced into students’ HPE course’s grade book, as well all other course-related survey data - including BMI values as calculated from self-reported weight and height values, and demographic data (age and sex) as part of this study. The data are from a four-semester period (Fall 2018 - Spring 2020) and were collated and de-identified by members of the institutional research team before being passed to us for analysis. This study does not include data from students who opted out (the default option), students with BMI values below 14.5 kg**·**m^-2^ or above 49.4 kg**·**m^-2^, and students whose age was below 16 or above 24. The final de-identified dataset is available as Open Data (CC0) via figshare.^32^ The protocol of this study was approved by the University’s Institutional Review Board.

### Measures

The University’s HPE faculty educate students about the importance of instituting and maintaining a physically active and healthy lifestyle, not only during their time in college but for the rest of their lives. To help encourage students to be physically active, the HPE department requires all students enroll in an HPE course each semester. All University HPE courses have a requirement of 150 active minutes per week and a step count goal equal to 10,000 steps per day; students use a Fitbit to help keep track of their steps and active minutes (as measured by time spent in various HR zones) in real time, which are then synced to their respective HPE course gradebooks. The summary statistics of the data can be found in Table 2, in the results section, for the following variables:

**Table 1.**
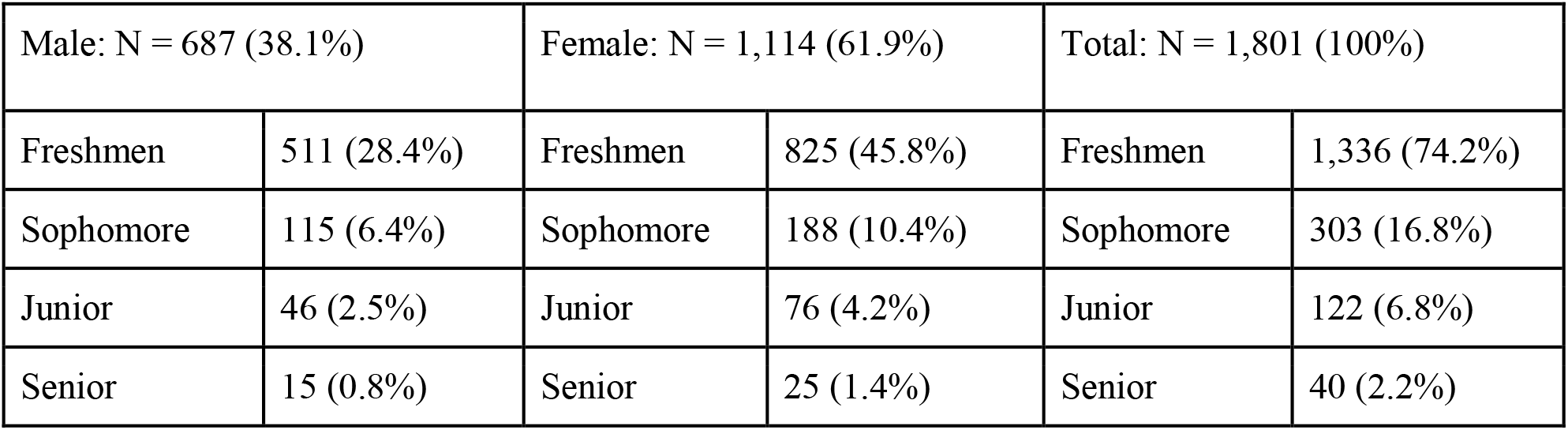
Summary of Study Participant Demographic Data

**Table 2.**
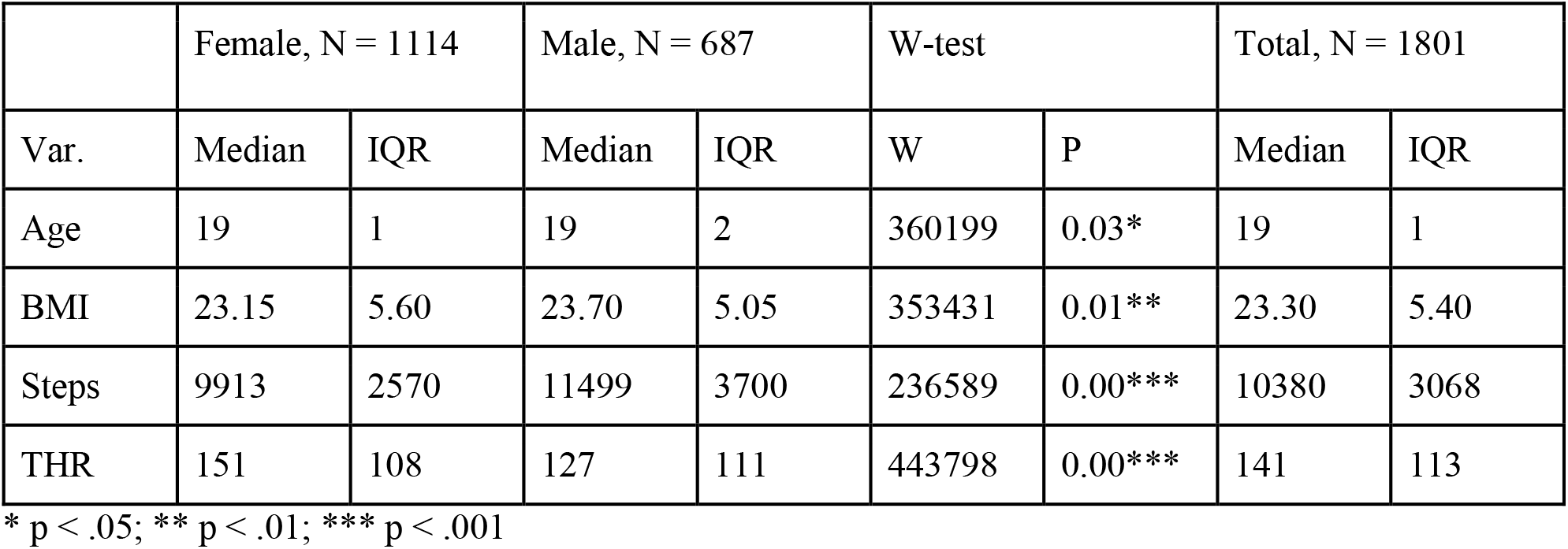
Summary statistics for the data used in this study, N = 1801.

#### Sex

The sex of the student was encoded as “0” female and “1” male. Data were collected from 1,114 females (62%) and 687 males (38%) – categorical variable.

#### Age

The age of the student, in years, approximately one week before each HPE course ended. This corresponds to the day the data are imported into our student records system for permanent storage. The study participants had a median age of 19 years and a mean age 19.02 years with a standard-deviation of 1.3 years – treated as a continuous variable.

#### BMI

The Body Mass Index (BMI) of study participants. BMI is taken and recorded during a lab and quiz questions test the students’ knowledge about it. The majority of instructors assist freshmen in accurately measuring and assessing their BMIs in these labs. After the freshman year, most instructors send the students to the BMI room to measure and self-report their own BMI values. The study participants had a median BMI of 23.3 kg**·**m^-2^ (IQR 5.4 kg**·**m^-2^) and a mean BMI 24.6 kg**·**m^-2^ with a standard-deviation of 5.2 kg**·**m^-2^ – treated as a continuous variable.

#### Steps

A Fitbit-recorded measure equal to the average number of steps that study participants took per day as a measure of physical activity over the approximately 15-week long HPE course period. For all our students we have a step count goal of 10,000 steps per day and step counts are a small part (typically 10%) of the student’s overall HPE course grade. For students who opted into the study (over 95%), daily step counts were measured using Fitbit devices and automatically synced to the course gradebook – treated as a continuous variable.

#### THR

The sum of weekly FatBurn, Cardio, and Peak HR zone minutes as a measure of total/training heart workload – treated as a continuous variable:

- FatBurn. A Fitbit-recorded measure equal to the total number of “Fat Burn” minutes per week. Fat Burn minutes are accrued when the student’s HR is in the Fat Burn zone typically between 100 beats per minute and 140 beats per minute for students in this study.
- Cardio. A Fitbit-recorded measure equal to the total number of “Cardio” active minutes per week. The Cardio zone is typically between 140 beats per minute and 170 beats per minute for students in this study.
- Peak. A Fitbit-recorded measure equal to the total number of “Peak” activity minutes per week. The Peak zone, a high-intensity exercise zone, corresponds to a heart rate greater than 85% of maximum, typically greater than 170 beats per minute for students in this study.

### Statistical analysis

Data handling, processing, and analyses were conducted using R version 4.0.2.^33^ All factors were tested for normally using a Shapiro-Wilk test with none being found to be normal and therefore the variables are described using median as the measure of central tendency (average) and interquartile range instead of mean and standard deviation, see Table 2 below. Sex differences were tested for by using an unpaired two-samples Wilcoxon test, Pearson’s correlation was used to analyze the association between all studied parameters, and one-way analysis of variance (ANOVA) was used to test between competing models. The values p < .05 were considered statistically significant.

## Results

### Demographics

The age in years (Age), body mass index (BMI), average daily step count (Steps), and weekly minutes with a HR above 50% of maximum (220 - Age), which equates to approximately 100 BPM for the participants of this study (THR), were collected from 1801 primarily first- and second-year university students enrolled in a required HPE course.

The student body at our institution is diverse. The demographics of the study participants are presented in Table 1 and the summary statistics for the measures used in this study can be found in Table 2, where THR (median = 141.3 min) consists primarily of Fat Burn zone minutes (median = 132.9 min) followed by Cardio minutes (median = 4.7 min) and then Peak minutes (median = 0.8 min).

### Sex Differences

BMI, Steps, and THR were each tested to see if there was a difference in values for men and women by using an unpaired two-samples Wilcoxon test with significant differences (p < 0.05) found for BMI (with men having a slightly higher average BMI value), Steps (with men taking an average of 1,586 more steps per day than women), and THR (with women spending an average of 24 additional minutes with heart rates above 100 BPM), see Table 2 and Figure 1.

**Figure 1.**
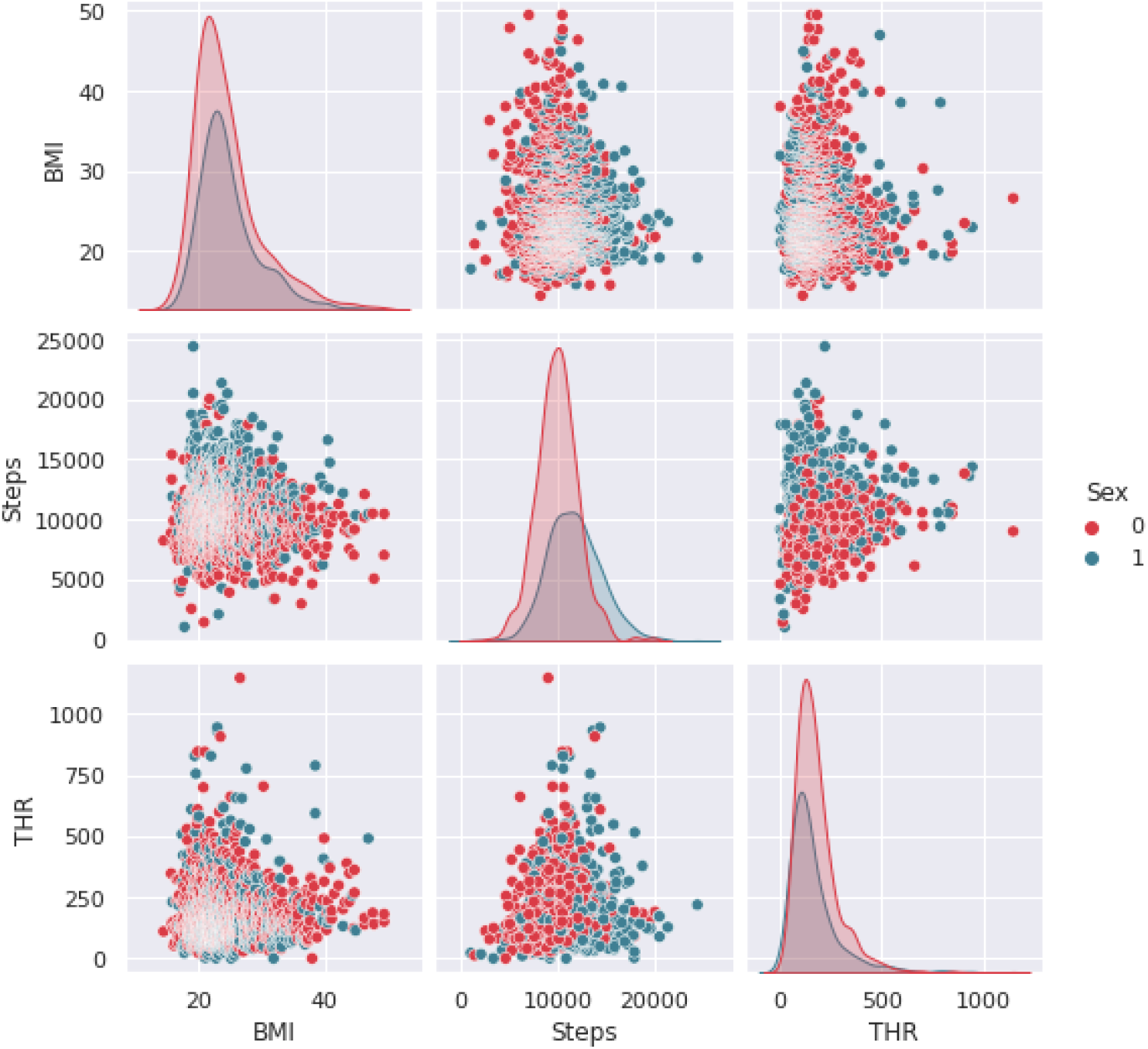
Steps and THR by Sex. Females and males have roughly the same BMI distributions (top left). Females have higher THR minutes than males (bottom right) but they have lower and less disperse steps per day than their male counterparts (middle).

### Correlations

The correlation matrix (Table 3), shows several significant correlations between the various variables in this study.

**Table 3.** Correlation matrix for study variables.

| Variable | BMI | Steps | THR | Sex | Age |
| --- | --- | --- | --- | --- | --- |
| BMI | 1.00 | -0.06* | 0.04<sup>•</sup> | 0.03 | 0.09*** |
| Steps |  | 1.00 | 0.06** | 0.33*** | -0.07** |
| HR |  |  | 1.00 | -0.06* | -0.01 |
| Sex |  |  |  | 1.00 | 0.07** |
| Age |  |  |  |  | 1.00 |

<sup>•</sup> $p < 0.1$ ; \* $p < .05$ ; \*\* $p < .01$ ; \*\*\* $p < .001$

### Physical Activity and BMI

There is a significant difference between men and women and their respective level of physical activity as measured by the average number of steps taken per day, with men on average taking 1,586 more steps per day than women (r = .33, p < .001). The negative correlation between BMI and Steps means that students with higher BMI values are generally less physically active (r = -.06, p < .05). However, when examining the plot of Steps vs. BMI, there is a clear non-linear relationship between the two, see Figure 2.

**Figure 2.**
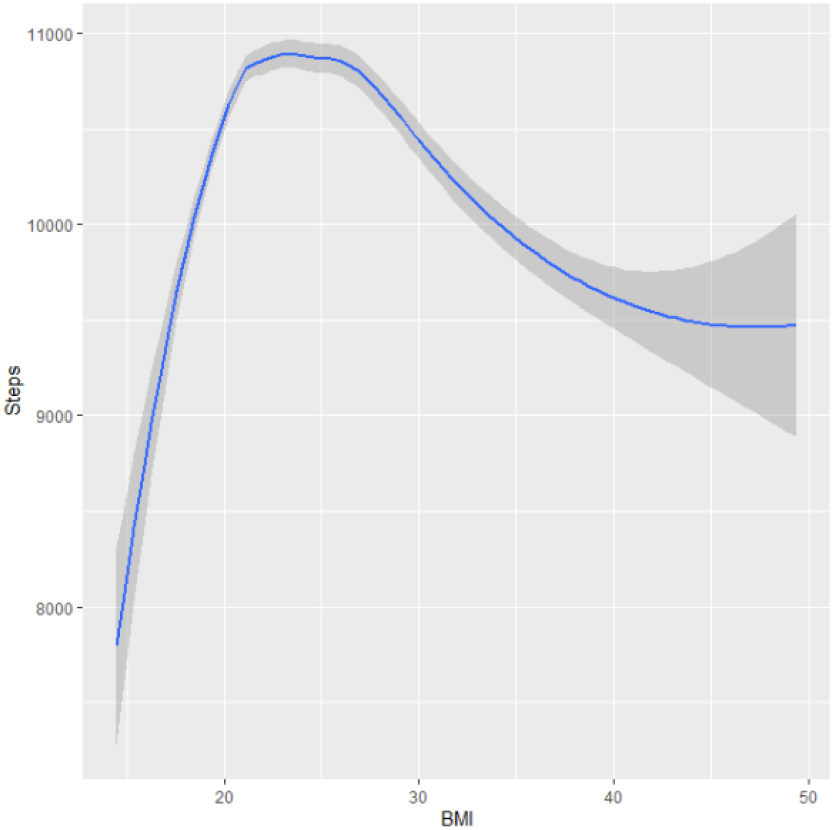
Steps vs. BMI. Both students with high BMI values (obese, BMI > 30 kg**·**m^-2^) and students with low BMI values (underweight, BMI < 18.5 kg**·**m^-2^) are less physically active, in general, than the other (normal and overweight, BMI range 18.5—30 kg**·**m^-2^) students in the study.

By performing an one-way analysis of variance F-test (ANOVA; F = 9.40, p = 0.002), we found that a quadratic model was a significantly better fit than a linear one, even when controlling for both Sex and Age (model R^2^ = .12, p < .001; p < .01 for all coefficients):

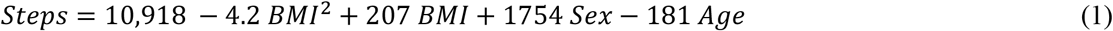

Note: the sex variable is assigned 0 for females and 1 for males.

### Training Heart Rate (THR) and Body Mass Index (BMI)

There is a significant difference between men and women and their respective heart load experience as measured by the total number of minutes per week with a high HR (THR; number of minutes per week with a HR >∼ 100 bpm), with women experiencing on average an extra 24 minutes per week of high HRs than men (r = -.06, p < .05). The positive correlation between BMI and THR means that students with higher BMI values generally experience more minutes per week with high HRs (r = .04, p < .1). By examining the plot of THR vs. BMI, we see that the relationship between THR and BMI, just like the relationship between Steps and BMI, is non-linear, see Figure 3.

**Figure 3.**
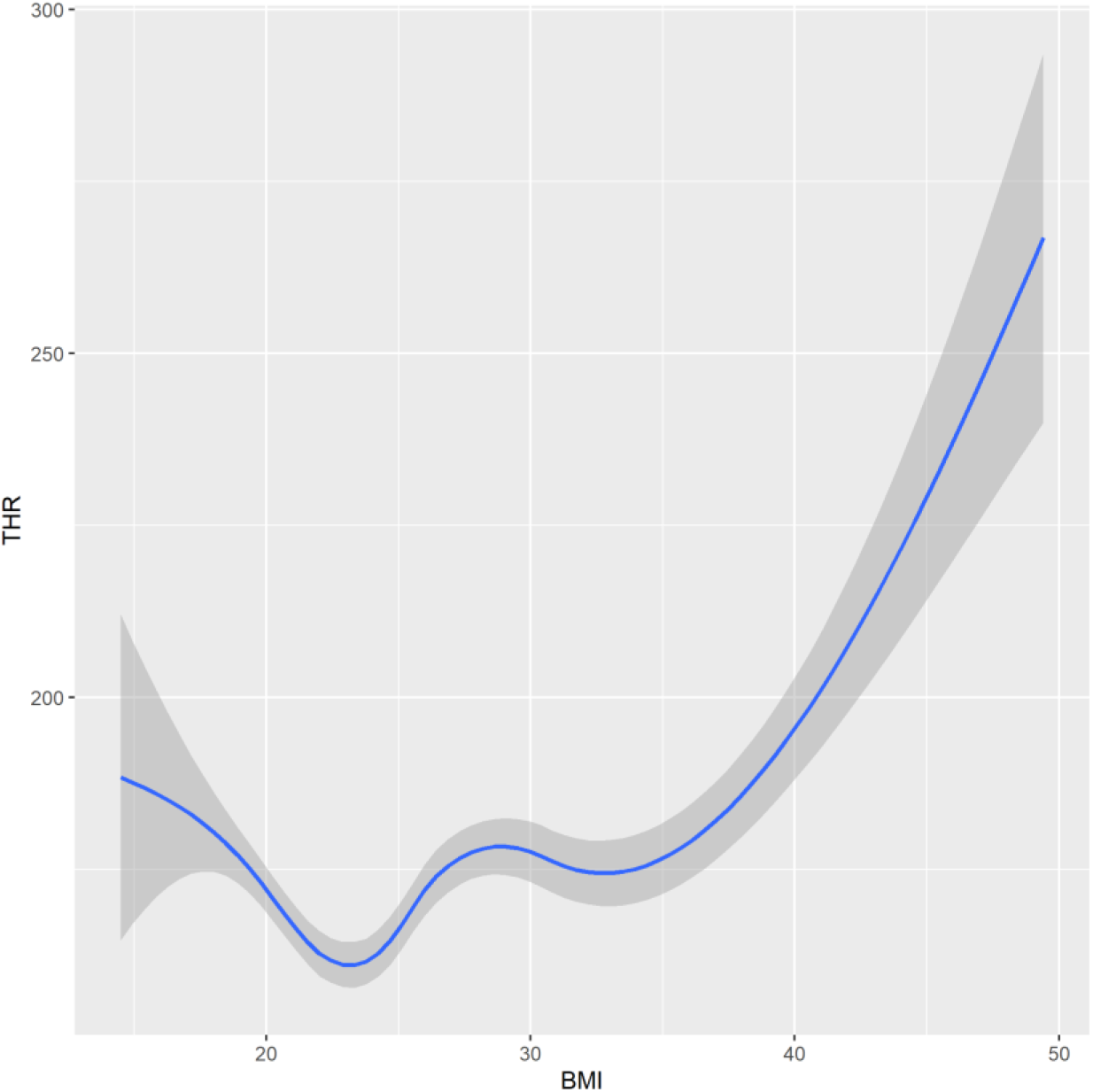
THR vs. BMI. Both students with high BMI values (obese, BMI > 30 kg**·**m^-2^) and students with low BMI values (underweight, BMI < 18.5 kg**·**m^-2^) experience more time with high heart rates (THR), in general, than the other (normal and overweight, BMI range 18.5—30 kg**·**m^-2^) students in the study.

By performing a one-way analysis of variance test (ANOVA; F = 6.20, p = 0.013), we found that a quadratic model was a significantly better fit than a linear one, even when controlling for Sex and daily physical activity as measured by Steps (model R^2^ = .02, p < .001; p < .05 for all coefficients):

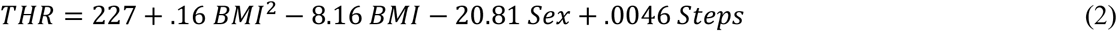

In order to compare the graphical outputs produced by R (Figure 2 and 3), both traces were plotted using the same x-axis (BMI) and presented on Figure 4. In this representation, it is clear that the BMI value corresponding to the minimum value of THR coincides with the BMI value for the maximum of Steps. It is observed where the concavity changes may indicate a change in the health zone for any normal individual. It is observed that changes in concavity and/or local minima of the model happen at BMI = 18 kg**·**m^-2^ and BMI = 28 kg**·**m^-2^, which match closely to the range of healthy weight reported by the CDC (18 kg**·**m^-2^ < BMI < 25 kg**·**m^-2^).

**Figure 4.**
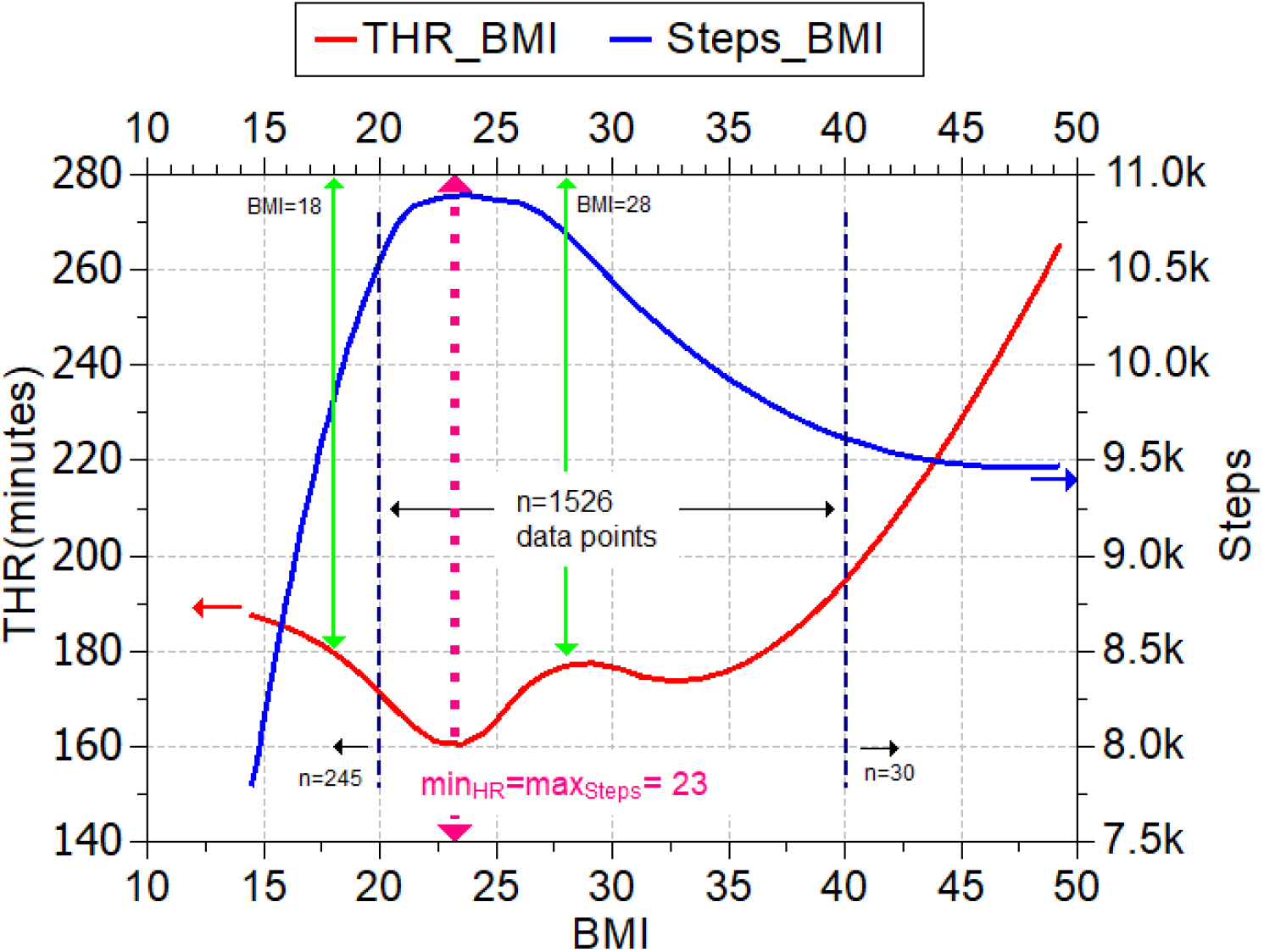
THR and Steps models plotted on the same axis. Minimum THR minutes and maximum number of Steps occur at approximately the same BMI value 23 kg·m^-2^.

In addition, we indicate on Figure 4 the number of data points that are in each zone of our model. For BMI > 20 kg**·**m^-2^ and BMI < 40 kg**·**m^-2^ there 1,526 individuals. For BMI < 20 kg**·**m^-2^, we have 245 individuals and for BMI > 40 kg**·**m^-2^ we have only 30 individuals. The increase in the error band (grey area) for our model, observed on Figure 2 and 3, is due to fewer data points being available in these extreme regions.

## Discussion

The purpose of this study was to correlate the heart workload profile of college students with their level of physical activity, sex, and BMI. Age in years, BMI, average daily step count, and weekly minutes with a HR above 50% of maximum (220 - age), which equates to approximately 100 bpm, were collected and analyzed from 1,801 primarily first and second year university students enrolled in a required HPE course.

Findings indicate that men took an average of 1,586 more steps per day than women, confirming what we observed in a previous study,^34^ and that women spent an average of 24 additional minutes per day with heart rates above 100 bpm (See Table 2 and Figure 1) even though they were, on the whole, less physically active. A significant negative correlation was found between BMI and steps meaning that students with higher BMI values (BMI > 30 kg**·**m^-2^) were generally less physically active. In addition, as previously conjectured by our HPE faculty, the data also showed that students with low BMI values (BMI < 18.5 kg**·**m^-2^) were less physically active than normal or overweight individuals (BMI range of 18.5-30 kg**·**m^-2^). This confirmed what our HPE faculty have observed qualitatively for decades and conforms recent results from Cho, Lee, & Kim.^21^ Thus it is not surprising that the relationship between Steps and BMI is better described by a quadratic model rather than linear one, see Figure 2.

When exploring the relationship between THR and BMI, a significant positive correlation was found which indicated that students with high BMI values (obese, BMI > 30 kg**·**m^-2^) experienced more time with a high HR, in general, than individuals with BMIs ranging between 18.5-30.0 kg**·**m^-2^. Similar to the results for Steps, students with low BMI values (underweight, BMI < 18.5 kg**·**m^-2^) also spent more time with high heart rates. This finding was interesting because one would not normally expect a high HR to be associated with low BMI values, especially when coupled with the fact that low BMI scores were also associated with low step counts. It seems that individuals with BMI scores below 18.5 kg**·**m^-2^ are much less physically fit (i.e. have a heart unused to physical activity) than their peers who were of normal weight or overweight. In fact, their results more closely resembled individuals with BMI scores over 30 kg**·**m^-2^.

Our research findings support those found in recent studies. For example, a study by Park, Lee and Han^17^ found that being underweight with a BMI less than 18.5 kg**·**m^-2^ increases one’s risk for cardiovascular disease, specifically stroke and heart attack, and more prominent in those under the age of 40. This may be attributed to the decline in body muscle mass as well as a possible relationship to poor nutritional status among underweight individuals. Similarly, a longitudinal study of more than 1 million women done by Canoy^15^ showed that a high BMI contributes to coronary heart disease and mortality risk as well as finding a low BMI was also associated with that same risk of disease and mortality. A study by Roy and McCrory^20^ also supported that those who were sedentary had higher max HR levels. This aligns with the low steps count and lack of physical fitness we observed due to the sedentary activity of our study participants with low BMI values.

### Limitations

The fact that THR and Steps were part of the students’ grades may have provided more incentive to be active than college students who did not have such importance placed on Fitbit data. This leads to the possibility for Steps and THR inflation by students who shook their Fitbit to collect steps or used other sources of heat to create a false THR reading rather than engaging in activity, though our previous studies have shown that the influence of Fitbit stacking is minimal.^34^

The correlations between the various variables, while significant, are, except for the correlation between the variables Steps and Sex, fairly small and therefore the results should be read accordingly, see Table 3.

Other limitations include the convenience nature of our study without a control group, the inability to control for resting heart rate values (something we’ve identified for a future study), and that BMI does not account for the muscle mass of certain individuals with high BMI scores.

## Conclusions

The findings of this study suggest that having a healthy BMI and physical activity level affects daily heart workload in a positive way and that individuals with either very low or very high BMI values have greater heart workloads than their normal weight peers. In addition, individuals with very low or very high BMI also take fewer steps per day resulting in a more sedentary lifestyle. Given the positive relationship between physical inactivity and risk of CVD, these findings indicate that both low and high BMI groups with corresponding lower steps and higher THR could be at higher risk of developing CVD than normal weight individuals.

The opportunity to promote healthy behavior lifestyles through increased physical activity is evident and ongoing for college students to prevent increases in weight gain, increased BMI, and cardiovascular risks throughout the college years and beyond. The fact that people with higher BMI had a higher heart rate was not surprising, due to their heart carrying a heavier load. One key finding from this study is that this research showed those with unhealthily low BMIs can more easily experience high heart rates than those in the normal, healthy BMI range due to their lack of fitness caused by a physically inactive lifestyle.

